# Non-invasive eDNA reveals the ecological and genetic status of the Western Capercaillie (*Tetrao urogallus aquitanicus*) in the Eastern Pyrenees

**DOI:** 10.1101/2025.01.14.632749

**Authors:** Pauline Buso, Céleste Paris, Manon Canal, Morgane Tassaint, Sylvain Daniello, Anaïs Gibert, Rupert Palme, Sabine Macho-Maschler, Christel Llauro, Gwenaël Jacob, Valérie Delorme-Hinoux, Joris A. M. Bertrand, Olivier Rey

## Abstract

The Anthropocene era is expected to bring about significant biodiversity and habitat loss for many species. These geographical changes, whether driven by climatic or anthropogenic factors, are likely to lead to considerable alterations in population size, structure, and genetic diversity. Monitoring natural populations is therefore essential to assess these impacts and enable informed conservation strategies for threatened species. The Western Capercaillie (*Tetrao urogallus*, L. 1758) has a widespread distribution in Boreal forests but fragmented in mountainous regions of the Palearctic, and is locally threatened by climate change, habitat destruction, and human disturbance. Our study focused on the eastern population of the subspecies *T. u. aquitanicus,* which is endemic to the Pyrenees mountains. The monitoring of this population has relied on direct methods and no genetic information had been generated so far. Here, we conducted a molecular study based on 229 non-invasive samples (faeces) to assess the ecological and genetic status of local population in the Catalan Nature Reserves of the Pyrénées-Orientales (Occitanie region, France). At the individual level, we assessed multi-locus genotypes, sexing, levels of inbreeding, stress level (Fecal Corticosterone Metabolites; FCMs) and diet. At the population level, we assessed sex ratio, genetic diversity and structure. We identified 62 individuals with a balanced sex ratio and estimated a census size of 79 individuals [95%CI = 68–92] in the study area. Genetic diversity was low and suggested significant inbreeding levels. FCM levels were lower in birds of areas considered as disturbed by humans and metabarcoding approach indicated a geographical structuring of diet composition at the reserve scale, with individuals exhibiting feeding behavior upon only one or few plant species. Our estimate of population census size was higher with figures assessed from lek counts, and the genetic approach provided additional insights on this population, establishing a baseline that will support conservation management plans.

## INTRODUCTION

Climate change and anthropogenic pressures are identified as the primary drivers of biodiversity loss (Ceballos *et al.,* 2015, 2020). These two interrelated factors result in habitat loss and fragmentation, which in turn gives rise to the isolation and local differentiation of populations. In mountainous regions, species are confronted with a dual challenge: the inherent difficulties posed by the topography, and the accelerating effects of climate change, which periodically push them to the limits of their survival and reproduction capacities. Many cool-climate species have already shifted their range to higher altitudes - often the last areas that meet their ecological needs (McCarty, 2001; Parmesan, 2006; Walther *et al*., 2005; Willis & Whittaker, 2000; Yousefi *et al*., 2015). Moreover, the rapid expansion of human infrastructure for outdoor recreation, livestock grazing, and the timber industry has exacerbated habitat destruction and fragmentation. These activities have reduced habitat connectivity for many species, impacting nearly 60% of mountainous regions by the late 2010s (Elsen *et al*., 2020; Nogués-Bravo *et al*., 2007; Rixen & Rolando, 2013; Schmeller *et al*., 2022; Wilson, 2016). Gene flow is a crucial factor influencing the spatial distribution of alleles and genetic diversity within a population, which partly determines its adaptive potential. However, it remains contingent upon the distance between populations, the dispersal capacities of species, and geographical relief. The lack of connectivity and gene flow between isolated populations results in an elevated risk of genetic erosion, inbreeding and ultimately local extinction (Bosse & Van Loon, 2022; Méndez *et al*., 2011; Mullu, 2016; Templeton *et al*., 1990). Therefore, it is crucial to obtain accurate estimates of the distribution, population dynamics, and genetic status of isolated populations at a fine geographic scale to establish conservation priorities and effective management strategies.

Investigating the ecological and genetic status of isolated populations is even more crucial in mountain areas as altitudinal gradients have created diverse ecosystems with high species richness and conservation value (Becker *et al*., 2007; Myers *et al*., 2000; Steinwandter & Seeber, 2023). Mountain Galliformes are cool-climate species that found refuge at northern latitudes and mountaintop habitats and are particularly threatened by current environmental changes and human activities (Chamberlain *et al*., 2016; McGowan, Owens & Grainger, 2012). Among them, the Capercaillie (*Tetrao urogallus*, L. 1758) is widespread in the contiguous Boreal forests from Siberia to Fennoscandia, and also occurs, albeit in smaller and more fragmented habitats in the mountain ranges of Central and Western Europe, including the Balkans, the Carpathians, the Alps, the Jura, the Pyrenees and the Cantabrian mountains (Duriez *et al.,* 2007; Leclercq, Bernard & Ménoni, 2018). In the Western part of its distribution, the species is usually associated with old-grown high-altitude beech and conifer forests (Suter, Graf, & Hess, 2002; Pakkala *et al.,* 2003). The Capercaillie is classified as “Least Concern” at a global scale by the IUCN (BirdLife, 2016), but several populations are locally threatened and the species is now legally protected in several Western and Central European countries (Storch, 2007). Since the 1990’s, the species’ decline has been mostly associated with destruction of forest habitat and intensive timber industry, excessive hunting, recreative outdoor activities, collisions with cables and fences, increased densities of meso-predators and consequences of climate change (Alba *et al*., 2022; Mikoláš *et al*., 2017; Moss *et al*., 2001; Segelbacher *et al*., 2008). In winter, the main source of food is pine needles, resulting in a low energy balance for the birds. Human disturbance can cause birds to flee, but the low energy balance at this time of year can be fatal and can limits, or even prevents, the likelihood of breeding encounters in spring (Thiel *et al.,* 2011; Rixen & Rolando, 2013; Formenti *et al.,* 2015). Simultaneously, climate warming may induce a shift in plant phenology (Inouye, 2022) and a mismatch between the peaks of resource availability, in early-spring, and peak of energetic demands by chicks in late-spring (Moss *et al.,* 2001).

The development of molecular techniques has enabled the use of various types of material, such as hairs, feathers, and faeces, deposited by wild individuals in the field as sources of DNA to conduct genetic studies (Taberlet & Luikart, 1999; Segelbacher, 2002). Combining non-invasive samples and molecular methods plays a pivotal role in the analysis and monitoring of threatened populations as it provides valuable support for conservation and recovery efforts (Leroy *et al.,* 2018; Supple & Shapiro 2018). In particular, obtaining genotypes from such non-invasive sampling allows estimating essential biodiversity variables (EBVs) including genetic diversity, genetic structure, inbreeding rates and effective population size, which are keystone metrics to harmonize and interpret biodiversity estimates from diverse sources (Hoban *et al.,* 2022). These EBVs allow to assess population viability (Scott *et al.,* 2020), identify populations with the greatest conservation need and potential sources of individuals for translocation programs and genetic rescue (Kyriazis *et al*., 2021; Supple & Shapiro, 2018; Van Der Valk *et al*., 2019; Whiteley *et al*., 2015).

Several mitochondrial and microsatellite markers, among others, have already been developed for capercaillie (Jacob *et al*., 2010; Lucchini *et al*., 2001; Segelbacher *et al*., 2000), enabling various studies to be carried out on genetic diversity, habitat fragmentation and phylogeny (Cayuela *et al*., 2019, 2021; Jacob *et al*., 2010; Lucchini *et al*., 2001; Rutkowski *et al*., 2017; Segelbacher *et al*., 2000, 2003, 2008). For instance, previous research has highlighted that capercaillie populations in boreal and central Europe suffer from low genetic diversity and inbreeding, raising significant concerns about their conservation status (Cayuela *et al*., 2019, 2021; Segelbacher *et al*., 2003). Indeed, capercaillie demonstrate restricted natal dispersal distances, which reduces gene flow among neighboring populations (Storch & Segelbacher, 2000). This restricted dispersal, combined with the philopatric behavior of males, likely exacerbates inbreeding within local populations. As in other grouse species, capercaillie reproductive behavior is centered around leks (i.e specific locations where males gather to perform courtship displays and attract females; Höglund & Alatalo, 1995). Notably, grouse species exhibit sex-biased dispersal patterns, with females being the primary dispersing sex, while males tend to establish territories close to their natal areas (Dunn & Braun, 1985; Small & Rusch, 1989). Outside of the breeding season, particularly during mid-winter, capercaillie from nearby leks form mixed-sex winter flocks, where they forage and roost together. When the breeding season begins, males return to the leks, which females visit for mating purposes (Höglund & Alatalo, 1995). Females disperse from the winter groups of their natal areas before their first breeding season, while males typically remain within their natal winter groups (Willebrandt, 1988). Consequently, leks tend to be composed of males from the same winter groups, suggesting that males within a lek may often be close kin. This structure may lead to kin selection and further influence genetic diversity and structure within capercaillie populations (Cayuela *et al*., 2019; Höglund *et al*., 1999) and to some extent lead to inbreeding depression.

Additionally, the combination of non-invasive samples with Next Generation Sequencing approaches also provides the opportunity to gather supplementary ecological information about individuals such as their diet (Cabodevilla *et al*., 2021; Ruppert *et al*., 2019). Non-invasive samples deposited by wild organisms contain material other than DNA, such as stress hormones metabolites, that can be quantified to evaluate the physiological status of host organisms in response to various stressors (Palme, 2019). For instance, by measuring corticosterone metabolites (FCMs) in droppings, Thiel *et al.,* (2011) established a positive correlation between the practice of outdoor recreative activities during winter and the physiological stress response of *T. urogallus* in the central Alps, although evidence is lacking that this response is associated with decreased fitness (Moss *et al.,* 2014).

The subspecies *Tetrao urogallus aquitanicus* has been geographically isolated during the last Ice Age and is now endemic to the Pyrenees mountains (Segelbacher & Piertney, 2007). This taxon is locally protected by nature conservation laws, Nature Reserves, and by the Natura 2000 framework (Bal *et al.,* 2021). Census size of the local population is monitored annually by national organisms such as Nature Reserves, the French Office for Biodiversity (OFB) and the Observatory for Mountain Galliformes (OMG). In the Eastern Pyrenees mountains, the Catalan Nature Reserves Federation (FRNC) was created during the 1980s to confer local suitable habitat and conservation management plans for ten renewable years. A baseline census size of the population was estimated at 57 individuals from data collected during lek counts (i.e counts of males calling in the mating areas, known as ‘leks’) carried out in 2014 in all known capercaillie habitats (S. Danielo. pers. comm). However, due to potential biases associated with observer variability and with the behavior of birds during leks, this approach serves primarily to assess population trends - whether the population declined or increased - rather than providing accurate population estimates. Despite of the efforts engaged to monitor capercaillie in Catalan Nature Reserves, methodological difficulties and biases associated with lek counts highlighted the need for complementary approaches. Many ecological and evolutionary aspects of the local capercaillie population remain unexplored, and no genetic studies have been carried out so far in this local area.

The present study aims at supplementing existing data from direct and rather invasive studies to better understand the ecological status of *T. u. aquitanicus* in the Eastern Pyrenees. To do so, we combined microsatellite genotyping, genetic sexing, metabarcoding, and FCMs quantification based on non-invasive samples collected in four areas within FRNC in order to (1) estimate a minimum population size and sex ratio, (2) assess specific estimated EBVs for this population, (3) compare individual levels of circulating FCMs in zones differing in terms of human disturbance (low, medium, and high disturbance levels), and (4) determine variation in diet among individuals.

In light of the known decline of the species, current trends in habitat fragmentation and the effects of climate change on mountain ecosystems, we expected a small population size for *T. u. aquitanicus* in the Catalan Nature Reserves. In addition, we expected low genetic diversity and inbreeding, probably due to the combined effects of limited population size, short spatial dispersal and geographical isolation, which collectively restrict gene flow. It was also anticipated that the inbreeding coefficient is higher in males than in females due to sex-biased dispersal distances and male philopatry. Regarding stress level, we expected higher levels of FCMs in individuals living in areas where winter tourism and disturbance was more important. Given that this study took place in winter, when dietary diversity was limited, we also expected to observe a feeding pattern dominated by pine needles. Altogether, these data complement the results of previous monitoring campaigns and provide a baseline of the capercaillie’s ecological status in the study area for conservation strategies in the Catalan Nature Reserves. On a wider scale, this study will enhance knowledge and understanding of the ecological status of the capercaillie in the Pyrenees.

## MATERIALS AND METHODS

### Study area and field survey

The territory of the Catalan Nature Reserves spans an area of 93.52 km^2^ with diverse habitats, including high-altitude biomes such as willow stands, beech forests, and alpine meadows. Indirect evidence (footprints, isolated droppings, feathers) of the presence of capercaillie was sought in the field between January and March 2022, by surveying all the known historical wintering spots (n = 14) distributed into four areas: the Py (n = 6), Mantet (n = 6), Prats-De-Mollo-La-Preste (n = 1) Nature Reserves and the frontier between Py and Mantet Reserves (n = 1). Droppings were collected along vertical and horizontal transects across each of the 14 sites, following contour lines and focusing on trees with low branches, as capercaillies primarily perch on these during the wintering period (Schroth, 1991; Graf, Mathys, and Bollmann, 2009). A total of 229 droppings were collected (one dropping per GPS point, see Fig. 1; Py (n = 141), Mantet (n = 68), Prats-De-Mollo-La-Preste (n = 6); frontier Py-Mantet (n =14)) on the snow during daytime surveys, by limiting as much as possible any disturbances, and stored in collection tubes at -20°C until further processing. We collected the freshest droppings possible by observing the appearance of the droppings (liquid or solid) as well as their dryness. GPS coordinates and date of collection were recorded for each sample. A muscle sample from a male individual was kindly provided by Kevin Foulché (French Biodiversity Office, Orlu city, Ariège 09 France, 05/19/2021, SAGIR no. 145327, autopsied at LPL on 07/06/2021) and was used as reference genotype and positive control for molecular sexing and genotyping.

**Fig. 1:**
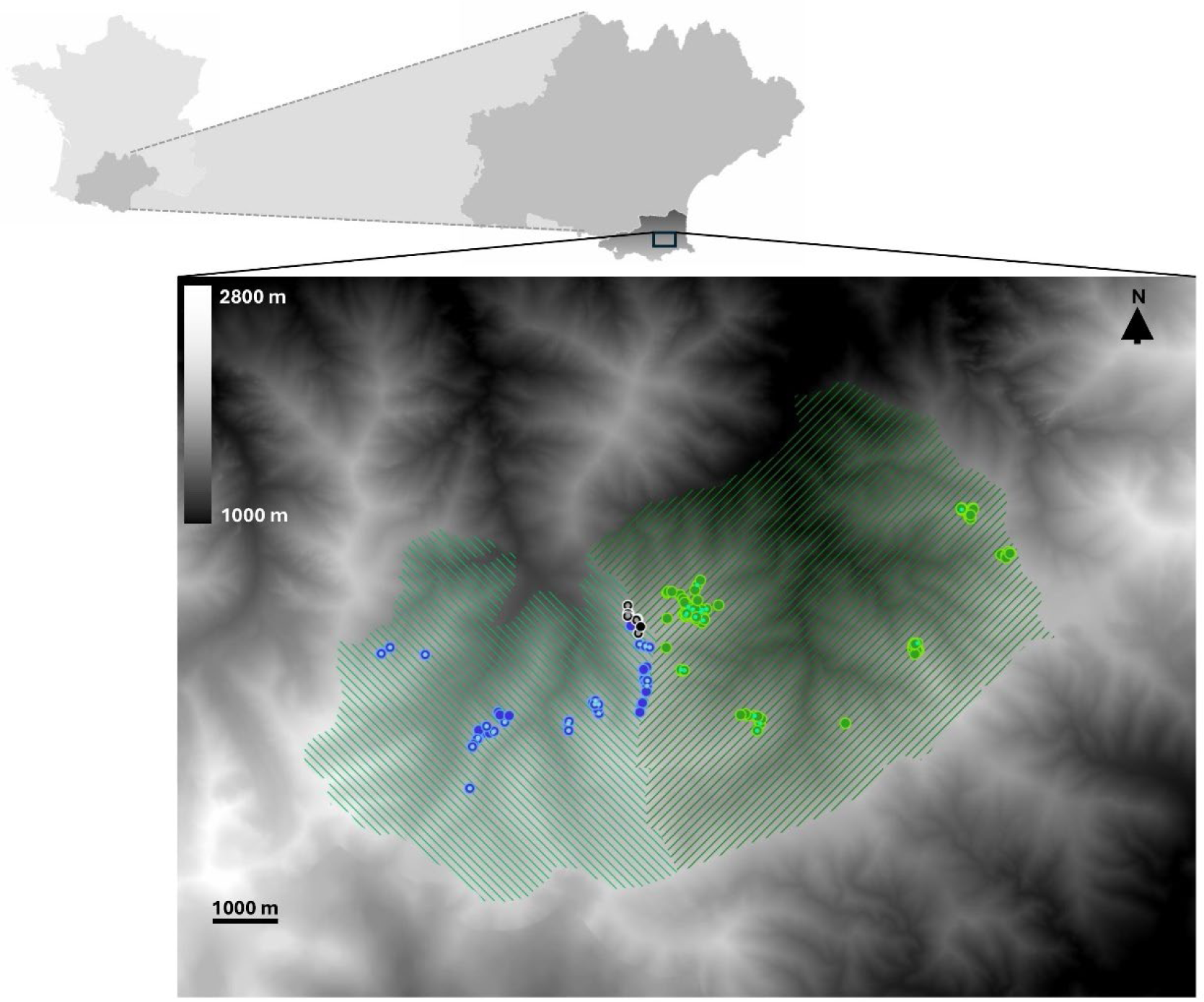
Sampling map of the 229 droppings of *Tetrao urogallus subsp. aquitanicus.* The droppings were collected in France, in the Occitanie region, more specifically in Eastern Pyrenees. The geographical limits of the reserves are visible by the zebra shapes (in light green for Mantet and in dark green for Py). The large dots correspond to the total sampling while the small dots correspond to the droppings used for diet analyses (in green, droppings sampled in the Py Nature Reserve; in blue, droppings sampled in the Mantet Reserve; in black droppings sampled at the frontier between Py and Mantet Reserves). The droppings surveyed in the Prats-De-Mollo-La-Preste Reserve are not visible on the map due to a GPS referencing issue. In order to protect the confidentiality of the data and in agreement with the Catalan Nature Reserves, the location of the sampling sites are not displayed.

### DNA extraction, molecular sexing and genotyping

Total genomic DNA was extracted from droppings using three commercial DNA extraction kits specific for stool samples due to delivery issues associated with the Covid-19 outbreak situation. We used EZNA® Stool DNA Kit (n=119 droppings, Omega Bio-Tek Inc, Norcross, USA) and QIAamp® DNA Stool Mini Kit (n=31 droppings, Qiagen, Hilden, Germany) following the manufacturer’s protocol for pathogen’s DNA extraction and extended the incubation time of the proteinase K at 70°C to 2 hours. We also used the NucleoSpin® DNA Stool Mini Kit for n=79 droppings (Macherey-Nagel GmbH & Co. KG, Düren, Germany) following the manufacturer’s protocol. DNA concentration for each extract was quantified using Nanodrop 2000. For extractions that did not pass the quality control, the droppings were extracted a second time (EZNA® Stool DNA Kit, n=18 droppings; QIAamp® DNA Stool Mini Kit, n=7 droppings).

Molecular sexing was achieved by PCR amplification of a fragment of the *CHD* region using primers P3’ and 1237L (Cayuela *et al.,* 2019). PCR was performed in 25 µL reactions containing 2 µL of DNA extract, 5X of GoTaq Buffer (Promega), 5 unit/µL of GoTaq® G2 DNA Polymerase (ref M7845 Promega), 2 µM of each dNTPs, 2 µM of each primer and RNase and DNA free water. PCR were programmed as follows: initial denaturation at 95 °C for 2 min, followed by 38 cycles of [denaturation at 95°C for 45 seconds, annealing at 55 °C for 2 min and extension at 72°C for 45 seconds] and a final extension at 72°C for 10 min. PCR products were run on a 3% agarose gel with 1X TAE. We assigned droppings to a female if we observed two alleles of size 244 bp (*CHD-Z*) and 269 bp (*CHD-W*), and to males if we observed only the *CHD-Z* allele. Droppings providing ambiguous results were amplified in up to three additional PCRs. After the third round of amplification, ambiguous genotypes were left unassigned.

We inferred multi-locus genotypes using eleven microsatellite markers (See Suppl. Table 1) and initially designed for Black grouse, *Tetrao tetrix* (BG10, BG15, BG18, BG20; Piertney and Höglund 2001), Chicken, *Gallus gallus* (ADL142, LEI098; Gibbs *et al.,* 1997), and Capercaillie, *Tetrao urogallus* (TuT1, TuT3, TuT4, sTuD1, sTuD3; Segelbacher *et al.,* 2000; Jacob *et al.,* 2010). Microsatellites were amplified into four PCR multiplexes using dye-labelled forward primers (Suppl. Table S1). PCRs were set in 12.5 µL reactions, containing 2 µL of DNA extract, 1X of Qiagen Multiplex PCR Master Mix, 0.2 µM of each primer and RNase/DNA free water. PCR program were programed as follows: an initial denaturation step at 95 °C for 15 min, 38 cycles of [denaturation at 94 °C for 30 sec, annealing at 57°C for 2 min, and extension at 72°C for 45 sec], and a final extension at 72°C for 15 min. PCR amplification products were checked on a 2% electrophoresis agarose gel and sent for fragment analysis (INRAE Gentyane genotyping platform, Clermont-Ferrand, France).

### Individual identification and population size estimation

Allele scoring was achieved using the GeneMarker® software (SoftGenetics, State College, USA; Holland and Parson, 2011). Droppings showing ambiguous profiles or non-amplification of alleles at one or more loci were amplified in a second round of PCR. Genotypes that were not unambiguously determined after the second round of PCR were coded as missing values.

We next reconstructed unique Multi Locus Genotypes (MLGs) based on pairwise dissimilarity distances between pairs of genotypes using the R package ‘poppr’ (Kamvar *et al.,* 2014). Droppings were assigned to a MLG if their genotypes differed by less than three alleles. MLGs were crossed with individual drop genetic sexing to ensure correct assignations. In cases where droppings from the same MLG, but differed by sex, all droppings of the MLG were turned into females to account for possible genotyping errors at the CHD loci. We graphically estimated the ability of our set of microsatellite loci to distinguish between unique MLGs using an accumulation curve using the R packages ‘poppr’ (Kamvar *et al.,* 2014) and ‘adegenet’ (Jombart, 2008).

To test for data reliability, we used Micro-Checker 2.2 (Van Oosterhout *et al.,* 2004) to check for the presence of allele dropout (ADO, i.e. one of the two alleles at a locus of a heterozygous individual is not detected), genotyping errors due to stuttering and null alleles of the markers. We used Genepop 4.7 (Raymond & Rousset, 1995; Rousset, 2008) to check for Linkage Disequilibrum (LD) between pairs of loci. We used CERVUS 3.0.3 (Kalinowski *et al*., 2007) to compute the probability that two individuals taken at random shared the same multi-locus genotype (i.e Probability of Identity (PI)) and accounting for population structuring and the presence of siblings (PI_sib_), and to assess the discriminatory power of the nine microsatellite markers selected (see results; Kalinowski *et al*., 2007; Mollet *et al*., 2015; Taberlet & Luikart, 1999; Waits *et al*., 2001).

We used the R package ‘CAPWIRE’ (Miller *et al*., 2005; Pennell *et al*., 2012) to estimate census size based on the frequency of recapture of each MLG. Sampling was conducted before the breeding season and in late winter, when mortality is reduced (Cayuela *et al*., 2021) and individuals still occupied their wintering ranges. We could therefore confidently assume that the population was closed during the sampling window. A capercaillie population in winter consists of a mixture of individuals differing in their spatial distribution and detection probability. Subadult and non-territorial individuals of both sexes show erratic behavior. The spatial distribution of the data has a minimal effect on the estimation of population size using CAPWIRE (Miller *et al*., 2005). Adult males occupy stable and non-overlapping home ranges distributed around the lek (Gjerde & Wegge, 1989). Females settle into dense forests stands, where they aggregate in small groups (Storch, 1995). Given the biology of the species, we can readily assume that individuals differ in deposition rate and, or detection probability. Hence, the assumption of even capture probability seems unrealistic. We therefore used TIRM model to estimate population size.

### Estimation of essential biodiversity variables (EBVs)

We inferred the level of genetic structuring of the population within our study area using a Principal Component Analysis (PCA), and determined the most likely number of genetic clusters contributing to the population (*k* = 1–8) using a clustering analysis (sNMF) with the ‘adegenet’ (Jombart, 2008) and ‘LEA’ R packages respectively (Frichot & François, 2015). Genetic diversity and deviation from panmixia across all loci was assessed by calculating the number of alleles (*A*), expected heterozygosity (*H*_E_), observed heterozygosity (*H*_O_), and *G*_IS_ using Genodive 3.06 Software (Meirmans, 2020). *P*-value were determined based on 9999 permutations. Analyses were carried out at population level, and by sex in order to estimate kinship level between males and between females.

### Stress response to human disturbance

Individual levels of stress hormone were assessed from a subset of 96 droppings, collected within the four study areas in sites classified *a priori* as “disturbed” (1 site in Mantet Reserve n=15; 1 site in Py Reserve, n=16), “mid-disturbed” (1 site in Mantet Reserve n=8; 1 site in Py Reserve, n=12; 1 site at the frontier Py-Mantet, n=11) and “undisturbed” (2 sites in Py Reserve, n=12 and n=16 respectively; 1 site in Prats-De-Mollo-La-Preste Reserve, n=6). Levels of human-induced disturbance were categorized by Nature Reserve agents, based on the intensity of outdoor activities, including hiking trails and off-piste skiing. We quantified the concentration of excreted FCMs using a competitive cortisone Enzyme Immuno-Assay (EIA), specifically targeting metabolites with a 3,11-dioxo structure (for details see Rettenbacher *et al.,* 2004). This EIA has been validated for assessing FCMs from droppings in Western Capercaillie (Thiel, Jenni-Eiermann, et Palme, 2005). Analyses were conducted at the University of Veterinary Medicine in Vienna, Austria (Vetmeduni), where FCMs were extracted in duplicate for each sample (i.e., replicates from two pieces of the same droppings by MLG) in accordance with the recommendations set forth by Palme *et al.,* (2013). Extracts were analysed in the cortisone EIA in duplicate. Coefficients of Variations (CVs) of duplicates were only a few %, but samples were repeated when CVs exceeded 10%. Interassay CV of a low and high concentration pool sample were 6.7% and 5.1%, respectively.

Difference in FCMs concentration was analysed using a series of Generalised Linear Mixed Models (GLMM) implemented in the ‘lme4’ package in R (Bates *et al*., 2015). As the likelihood of the different models tested could not be differentiated based on Akaike Information Criterion (difference in AIC < 2), an information theoretic model averaging approach was adopted using the ‘MuMIn’ package (Grueber *et al.,* 2011). The model implemented in MuMIn was chosen for its biological relevance. In our case, two parameters were considered as “fixed” effects, i.e. the area of sampling collection in respect with its expected level of human disturbance (“disturbed”, “mid-disturbed”, “undisturbed”) and the sex of the individual (male or female). The “biological replicate” was set as a random effect. The interaction effect between these two fixed variables was also taken into account. The resulting model was coded as follow: lmer(FCMs concentration ∼ Level of disturbance * Sex + (1|Replicates). We tested for and validated assumptions of normality, homoscedasticity, overdispersion of the model, and consistency between replicates.

Assessment of the bird’s diet:

In the Catalan Nature Reserves, the Western Capercaillie is exclusively found in forests dominated by mountain pine (*Pinus uncinata*), which are associated with other characteristic plants, trees, and shrubs of mountain ecosystems. These include species such as rowan (*Sorbus aucuparia*), rhododendron (*Rhododendron ferrugineum*), wild rose (*Rosa* spp.), raspberry (*Rubus* spp.), bilberry (*Vaccinium* spp.), heather (*Calluna vulgaris*), honeysuckle (*Lonicera caerulea*), woodrush (*Luzula* spp.), wavy hair-grass (*Deschampsia flexuosa*), vetch (*Lathyrus* spp.), bedstraw (*Galium* spp.), common bent (*Agrostis capillaris*), red fescue (*Festuca rubra*), Scots pine (*Pinus sylvestris*), juniper (*Juniperus* spp.), purging broom (*Cytisus oromediterraneus*), common speedwell (*Veronica officinalis*), bearberry (*Arctostaphylos uva-ursi*), pale toadflax (*Linaria repens*), and silver birch (*Betula pendula*), among others.

The three Nature Reserves differed in forest habitat composition. The Py reserve exhibited a high diversity of forest habitats. Mesophilic pinewoods on siliceous soils and acidophilic pinewoods dominated by mountain pine, often associated with common speedwell, covered 46% of the forest area. Pioneer formations, consisting of birch (*Betula spp*.), willow (*Salix spp*.), and hazel (*Corylus avelana*), accounted for 31%. Riparian forests, including alder (*Alnus spp*.) and elm (*Ulmus spp*.), covered 1%, while acidophilic beechwoods accounted for 8%. Subalpine fir forests with rhododendron made up 7%, and conifer plantations, such as Scots pine (*Pinus sylvestris*), Douglas fir (*Pseudotsuga menziesii*), and Norway spruce (*Picea abies*), covered 4%. Ravine forests, such as the hygrophilous lime forests of the Pyrenees, represented 3% of the forest cover. In the Mantet reserve, mesophilic and acidophilic mountain pinewoods with common speedwell dominated, covering 79% of the area. Subalpine fir forests with rhododendron covered 10%, while pioneer formations represented 10.8%. Beech and oak woods were marginal, covering only 0.2% of the area. Finally, the Prats-De-Mollo-La-Preste Reserve was primarily characterized by mountain pinewoods dominated by mountain pine. It also included Pyrenean-Cantabrian beechwoods, mountain and subalpine birch woods, conifer plantations, and pioneer formations, though the exact proportions of these habitats were not quantified (Danielo, pers.comm.).

In winter, the western capercaillie feed mainly on pine needles. We aim to determine whether we can potentially find other ingested plants by locally established birds despite the expected monospecific diet at this period of the year. We described individual diet following a metabarcoding approach based on a fragment of the chloroplastic *trnL* gene, amplified using the *trnL*-F (CGAAATCGGTAGACGCTACG), and *trnL*-R (GGGGATAGAGGGACTTGAAC) primers pair (Taberlet *et al.,* 2007) and Universal Illumina sequencing adapters.

Amplifications were performed in 25 µL reactions containing 2 µL of DNA extraction 5X of GoTaq Buffer (Promega), 5 unit/µL of GoTaq® G2 DNA Polymerase (ref M7845 Promega), 2 µM of each dNTPs, 2 µM of each primer and RNase and DNA free water. The PCR program consisted in a first denaturation step at 95 C° for 2 min, followed by 40 cycles of [denaturation at 95°C for 45 s, annealing at 52°C for 2 min and extension at 72 °C for 45 s] and a final extension at 72°C for 10 min. PCR products were run on 1% agarose gel to visually assess the PCR efficiency. Out of 229 droppings amplified, 49 showed clear amplification patterns and were sent for library preparation and 2x150 pb paired-end sequencing on a MiSeq 2000 (Illumina) at the BioEnvironment platform (University of Perpignan Via Domitia, France).

Sequencing reads were processed using the FROGS bioinformatic pipeline implemented in the Galaxy platform (Escudié *et al.,* 2018). After a preprocessing step, we clustered sequences into Operational Taxonomic Units (OTUs) using the swarm algorithm and an aggregation clustering distance of three. The resulting dataset was filtered out for chimeras using VSEARCH (Rognes *et al.,* 2016) and rare clusters (relative abundance < 5 x 10^-5^) were removed. Droppings showing less than 5000 reads were removed from the subsequent analyses. We assigned OTUs to taxa using NCBI Blast program (non-redundant nucleotide collection, updated in Oct. 2023). OTUs not assigned to *Plantae* taxa were excluded, as well as OTUs showing less than 97.5% identity with reference sequence. Since most OTUs were affiliated to several reference sequences from different species, we chose the best hit based on the Blast e-Value score and on the ‘Total score’ in the case of same e-Values. We also checked whether the assignations with similar e-Values and total score values belong to the same species or genus which was always de case. For sake of conservatism, we interpreted the resulting assignments at the taxonomical level at which all the best hists converged. The obtained raw OTU abundance matrix was rarefied based upon the smallest total abundance observed over droppings (n = 5746; see results).

Using the final OTU abundance dataset, we performed a Non-Metric Dimensional Scaling analysis (NMDS) based on pairwise Jaccard similarity indices to visualize and interpret diet composition similarity between droppings. A permutational multivariate analysis of variance (PERMANOVA; Anderson, 2017) was run to assess the effect of sampling area and individual sex on observed diet, accounting for interaction between these two variables. *P*-values were calculated based on 9999 permutations. The NMDS and the PERMANOVA were computed in R using the ‘Vegan’ package (Dixon, 2003). We also ran an indicator taxa analysis based on 9999 randomized permutations as implemented in the ‘indicspecies’ package (Cáceres & Legendre, 2009) to detect association between OTUs and study sites.

## RESULTS

### Reliability of the data

Out of the 229 collected droppings, 179 (78.2 %) were successfully genotyped and 203 (88.7 %) were unambiguously sexed. We did not observe obvious difference in amplification success between the three commercial kits used in our study. We found no evidence for possible allele drop out, null alleles or errors due to stuttering for most microsatellite markers. Only loci BG10 and TuT1 displayed an excess in homozygosity suggesting possible null alleles. No linkage disequilibrium was observed between pairs of loci except these two same markers BG10 and TuT1. Accordingly, subsequent analyses were conducted with the exclusion of these two markers. The nine remaining microsatellite markers were polymorphic, with a number of alleles ranging from two to four alleles, except for the marker LEI098, which was monomorphic (Table 1).

**Table 1.** Estimation of genetic diversity and deviation from panmixia across all loci, analyzed at the levels of the overall population, males, and females. The analysis includes calculations of the number of alleles (*A*), expected heterozygosity (*H*_E_), observed heterozygosity (*H*_O_), and the inbreeding coefficient (*G*_IS_). Significant estimates are highlighted in bold (*p-value* < 0.05).

| Global (*n*=62) |  |  |  |  | Males (*n*=28) |  |  |  | Females (*n*=33) |  |  |  |
| --- | --- | --- | --- | --- | --- | --- | --- | --- | --- | --- | --- | --- |
| Locus | *A* | *H<sub>O</sub>* | *H<sub>E</sub>* | *G<sub>IS</sub>* | *A* | *H<sub>O</sub>* | *H<sub>E</sub>* | *G<sub>IS</sub>* | *A* | *H<sub>O</sub>* | *H<sub>E</sub>* | *G<sub>IS</sub>* |
| LEI098 | 1 | 0.000 | 0.000 | *NA* | 1 | 0.000 | 0.000 | *NA* | 1 | 0.000 | 0.000 | *NA* |
| TuT3 | 3 | 0.097 | 0.123 | 0.216 | 3 | 0.143 | 0.201 | 0.289 | 2 | 0.030 | 0.030 | 0.000 |
| TuT4 | 2 | 0.419 | 0.401 | -0.045 | 2 | 0.536 | 0.429 | -0.250 | 2 | 0.333 | 0.389 | 0.144 |
| BG15 | 3 | 0.557 | 0.672 | **0.171** | 3 | 0.464 | 0.677 | **0.314** | 3 | 0.656 | 0.677 | 0.031 |
| BG18 | 3 | 0.452 | 0.424 | -0.065 | 3 | 0.429 | 0.456 | 0.060 | 3 | 0.455 | 0.395 | -0.151 |
| StuD3 | 3 | 0.393 | 0.415 | 0.051 | 3 | 0.407 | 0.447 | 0.089 | 3 | 0.394 | 0.403 | 0.022 |
| StuD1 | 3 | 0.267 | 0.305 | 0.126 | 3 | 0.259 | 0.383 | 0.323 | 2 | 0.281 | 0.245 | -0.148 |
| BG20 | 2 | 0.016 | 0.016 | -0.000 | 1 | 0.000 | 0.000 | *NA* | 2 | 0.030 | 0.030 | 0.000 |
| ADL142 | 4 | 0.311 | 0.389 | 0.199 | 4 | 0.296 | 0.430 | 0.311 | 3 | 0.333 | 0.369 | 0.097 |
| Overall | 2.667 | 0.279 | 0.305 | **0.085** | 2.556 | 0.282 | 0.336 | **0.162** | 2.333 | 0.279 | 0.282 | 0.010 |

### Estimated census size of the local Capercaillie population

The probability that two individuals shared the same multi-locus genotypes was low (PI = 0.004) even when considering population structuring and the presence of siblings (PI_sib_ = 0.0736). Our set of microsatellites provided enough power to distinguish individuals based on their multi-locus genotypes. Indeed, complementary analyses showed that a minimum number of seven loci was required to unambiguously assign genotypes to one of the MLGs (See Suppl. Fig. 3).

Pairwise dissimilarity analyses suggested the occurrence of 62 MLGs (see Suppl. Fig. 2), of which 33 were identified as females, 28 as males and one individual of unknown sex. Maximum likelihood estimator (MLE) using the two innate rate model (TIRM) was 79 individuals [95%CI = 68–92]. Relative difference in capture probability between types of individuals (harder *vs.* easier to capture, *α* = 4.0) confirmed our hypothesis of capture heterogeneity between individuals. MLEs by sex were 37 male individuals [95%CI = 30–47; *α* = 4.1], and 38 female individuals [95%CI = 33–47; *α* = 3.3]. The average capture frequency per individual was 2.89, with males exhibiting an average of 2.61 and females an average of 3.18.

### Essential biodiversity variables (EBVs)

In terms of genetic diversity and structure, the first two dimensions of the Principal Component Analysis explained respectively 13.4% (PC1) and 12.3% (PC2, sum = 25.7%) of the total genetic variance (Fig. 2). Overlapping distribution of individuals from the four reserves in the two dimensions of the PCA space based on their genotypes show no, or low levels of spatial genetic structuring at the scale of the study area. This is in line with the output of the clustering analysis (sNMF) which show that individual genotypes originated from a single gene pool (*k* = 1). At the population level, we observed low levels of heterozygosity (mean *H*_O_ = 0.279, mean *H*_E_ = 0.305), a deficit in heterozygotes and a significant positive inbreeding coefficient (*G*_IS_ = 0.085, *p*-value < 0.05). At the sex level, we found a mean *H*_O_ of 0.279, a mean *H*_E_ of 0.282, and a non-significant *G*_IS_ of 0.01 for females. For males, we observed a mean *H*_O_ of 0.282, a mean *H*_E_ of 0.336, and a significant positive *G*_IS_ of 0.162 (*p*-value < 0.05).

**Fig. 2:**
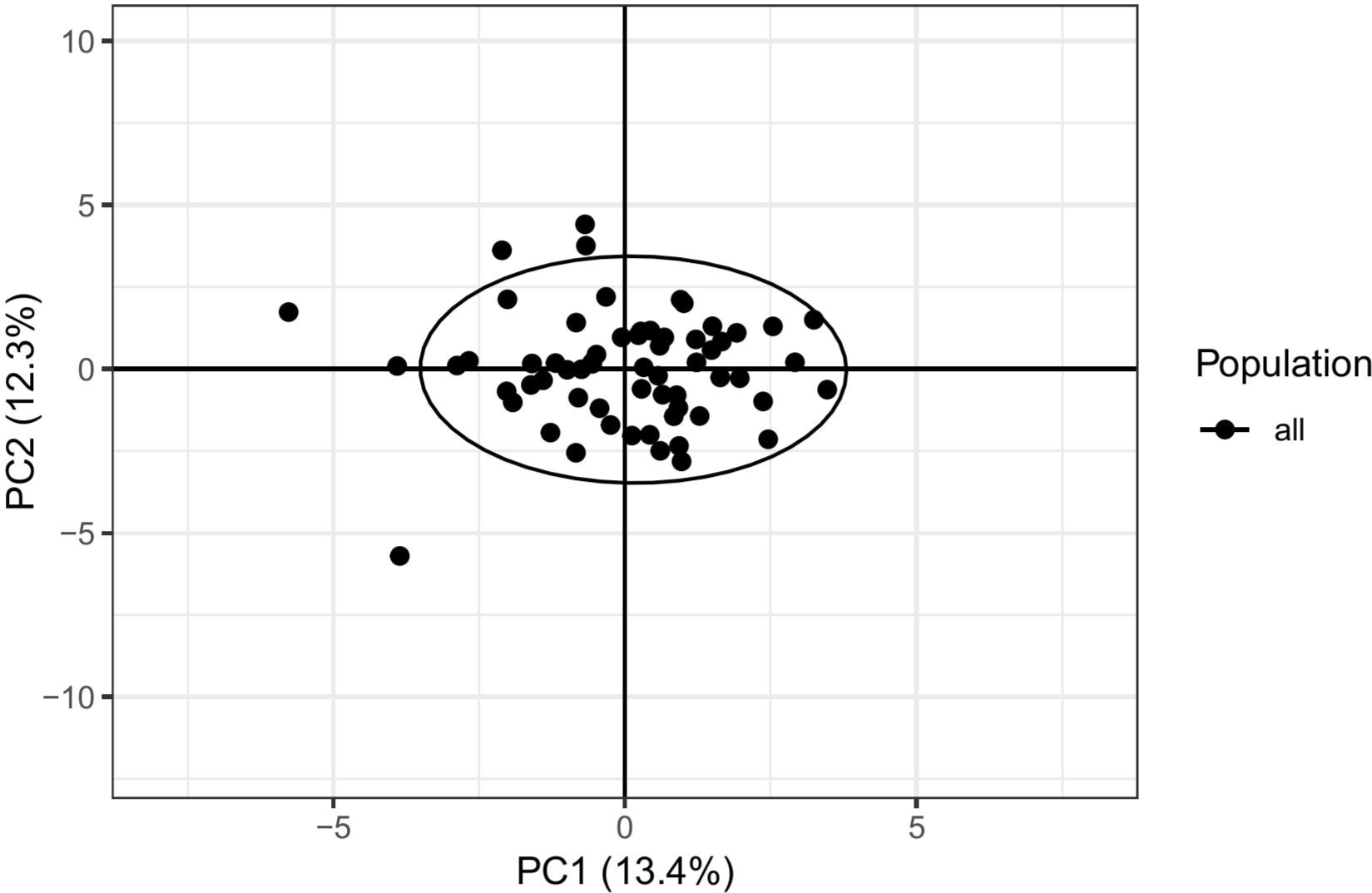
Principal Component Analysis (PCA) obtained from the 62 MLGs presents in the Catalan Nature Reserves. The first two principal components (PCs) jointly accounted for 25.7% of the total variance. The analysis shows that the capercaillies from the different reserves form a single entity of genetic structure.

### Individual levels of FCMs assessed from droppings

Regarding FCMs, the study site was the only variable that significantly impacted levels of FCMs (Fig. 3). *A priori* level of human disturbance was negatively correlated with individuals levels of FCMs (Estimate = -0.40; CI : [-0.597; -0.204] CI 95%). In other words, individuals had lower FCMs levels in areas *a priori* defined as disturbed (mean = 172 ng of FCMs/g of droppings) compared to the individuals at *a priori* undisturbed (mean = 335 ng of FCMs/g of droppings; Fig. 4). We neither observed difference in levels of FCMs between females and males (Estimate = -0.0408; CI : [-0.562; 0.481]), nor regarding the interaction between sex and disturbances levels (Estimate = -0.063; CI = [-0.460; 0.333]). Overall, our model explained 14% of total variance (adjusted R² = 0.139).

**Fig. 3:**
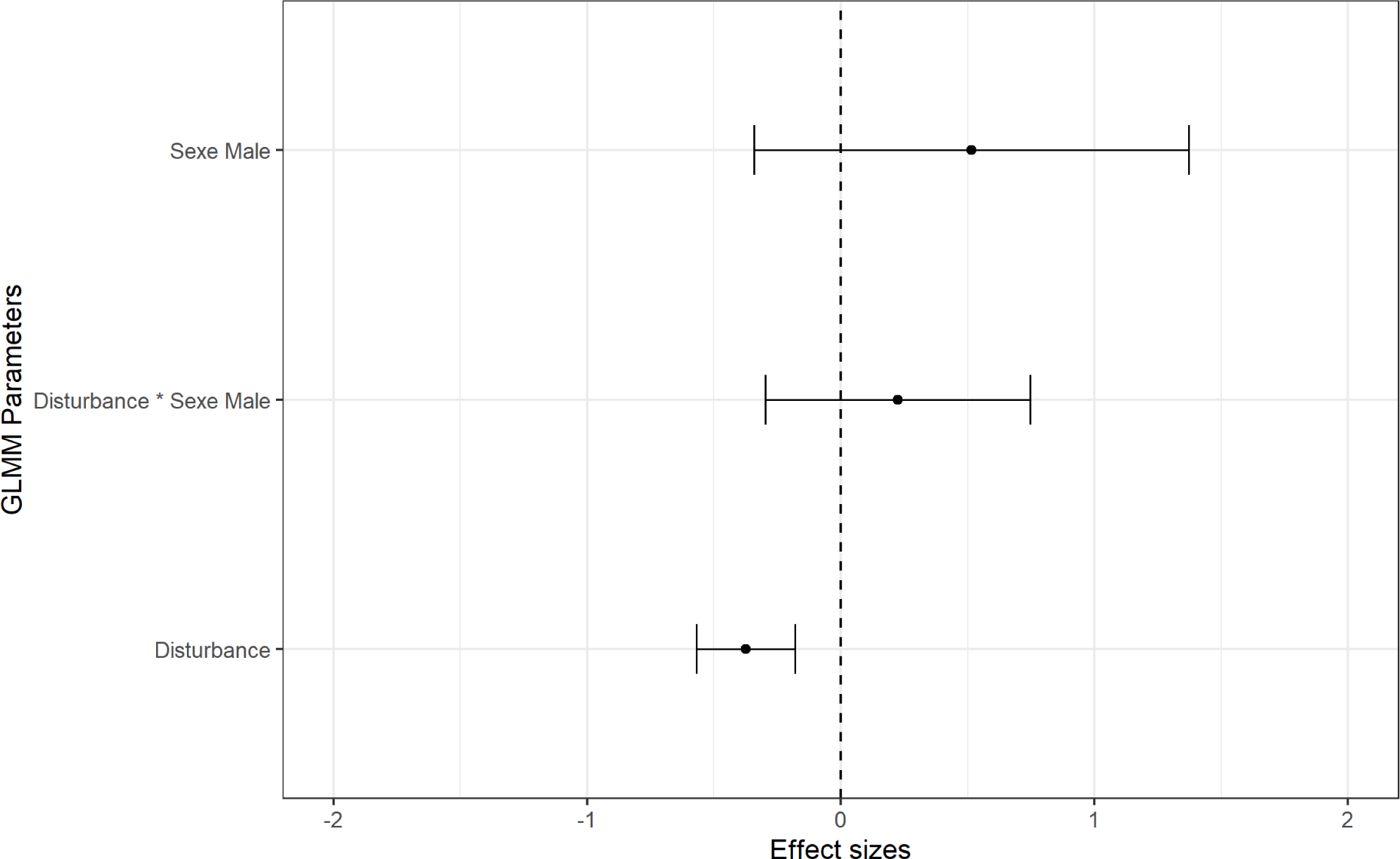
Estimates and CI values of the stress level by model averaging. Overall, 14% of the variance is explained by the model (R² adj = 0.139).

**Fig. 4:**
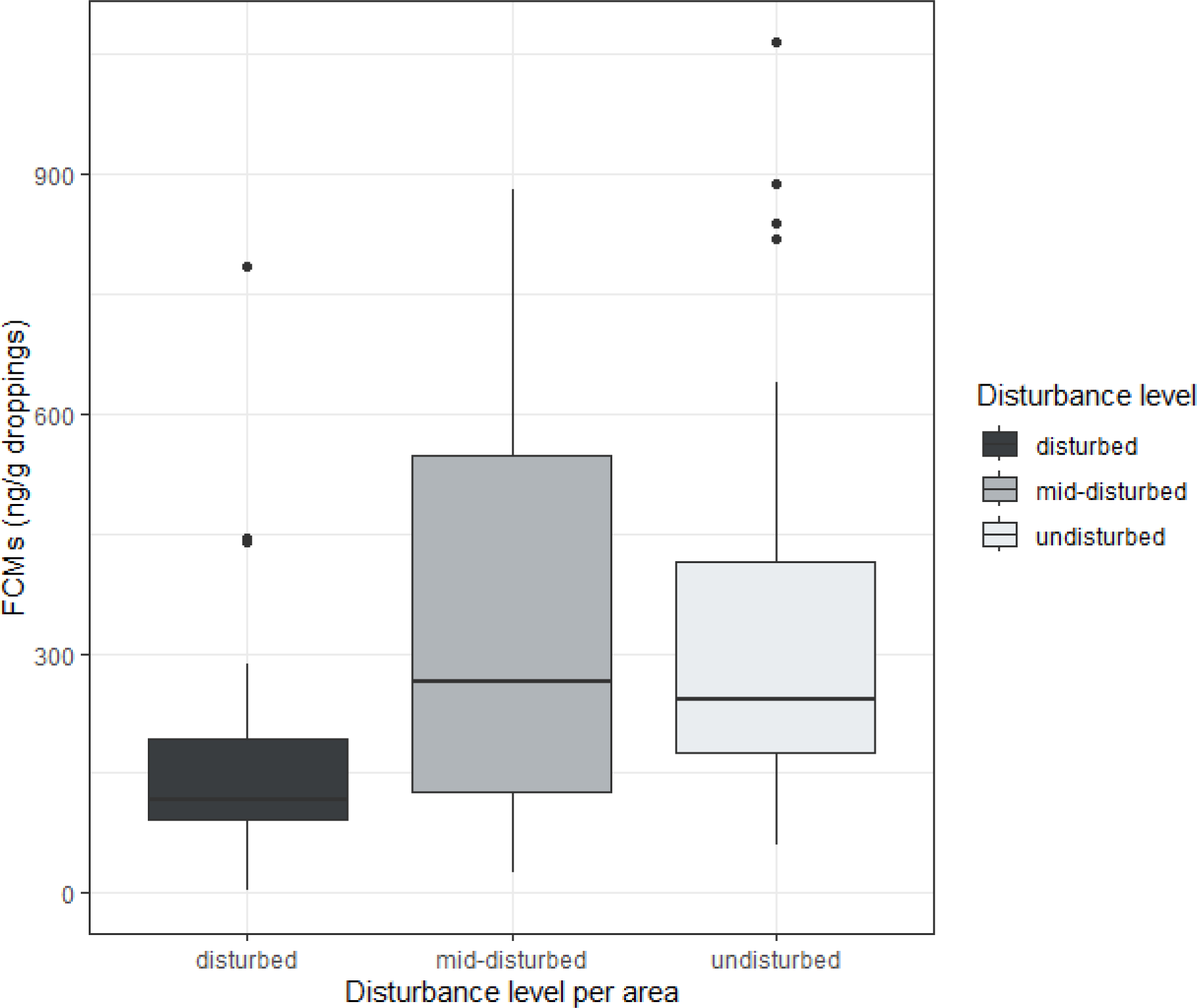
Boxplot of FCMs concentration (mean of two replicates (see Suppl. Fig. 5) in ng/g droppings) as a function of the GPS positions of the droppings and the estimated level of disturbance. The droppings from *a priori* “disturbed” sites show a trend of lower stress levels than droppings from *a priori* “mid-disturbed” and “undisturbed”.

Assessment of the bird’s diet:

Out of 229 droppings analyzed, 49 (21.4 %) showed enough amplification product on agarose gel and were sequenced, providing on average 86011 reads (range: 69–145785 reads).

After filtering and processing (FROGS pipeline), excluding OTUs not assigned to taxa within *Plantae* or showing less than 97.5 % identity with reference sequences, and excluding droppings with low amplification success (number or reads < 5000), we obtained a matrix of 29 OTUs and an average of 81827 reads per sample (range: 5796–141564) from 48 out of the 49 samples analysed. Three OTUs assigned to *Arabidopsis thaliana* and *Allium* sp. may have resulted from cross-contamination during the laboratory processing and were excluded. OTUs matching multiple species were assigned at the family level. Our final dataset thus comprised 26 OTUs belonging to 16 plant families. The number of OTUs detected within droppings ranged from 1 (12.2%) to 6 (8.2%) with most droppings containing 2 (38.8%) or 3 (26.5%) OTUs.

Based on the NMDS analysis (Fig. 5) and PERMANOVA analysis, the study site (Py, Mantet, Py-Mantet; Fig. 1) was the only variable explaining variation in individual diet (*F* = 2.75; *p*-value = 0.02). Neither the sex of capercaillie individual (*p*-value = 0.19) nor the interaction between the sampling area and sex (*p*-value = 0.07) contributed significantly. The OTU15 (*Betulaceae*) was exclusively found in the Py area and was identified as an indicator of study site (*p*-value = 0.017). The OTU18 (*Vaccinium* sp.) was almost exclusively found in Mantet and could be marginally considered as an indicator OTU (*p*-value = 0.076).

**Fig. 5:**
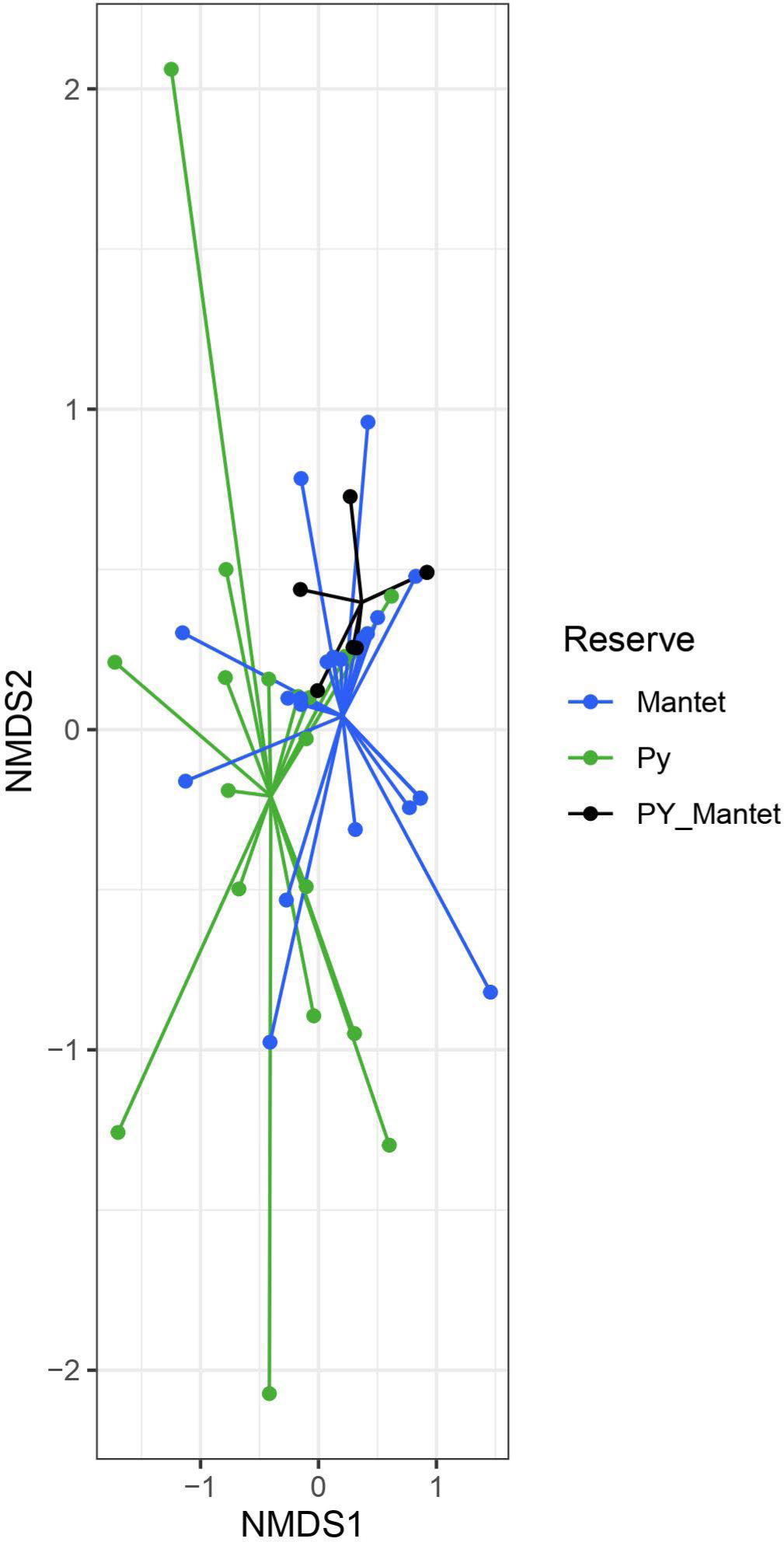
NMDS analysis showing difference in diet composition similarity between droppings among Py, Mantet and at the frontier of Py-Mantet Reserves.

## DISCUSSION

In the present study, we combined non-invasive sampling methods, molecular and analytic approaches to assess the ecological status of the Western Capercaillie population in the Catalan Nature Reserves. Our study provided an estimation of the census size and genetic characteristics of the population, as well as an evaluation of individual levels of stress-related hormones and diet selection. Our results provide a better understanding and knowledge of the ecological profile and genetic health of capercaillie in Catalan Nature Reserves, that could help guiding management plans at a local level.

### The local population of Western Capercaillie in Catalan Nature Reserves presents a low census size and a balanced sex ratio

The high level of genotyping success observed in our study confirmed the efficacy of combining non-invasive sampling, microsatellites genotyping and genetic sexing for the monitoring of populations, and especially for species that are difficult to observe like capercaillie. Non-invasive sampling, and especially droppings, generally contain minute amounts of highly degraded DNA, which increases the risk of missing some alleles of a heterozygote and overestimating of the proportion of homozygotes (Gagneux *et al.,* 1997; Taberlet, 1996). Our microsatellite control process pointed toward such a problem at only two among the 11 loci used, which were removed from the subsequent analyses. Despite the low polymorphism of the nine remaining loci, the probability of identity based on these loci event when accounting for possible sibship relationship between individuals were low, which provided strong support to clearly identify different individuals according to their genotypes. Moreover, the combination of seven loci was sufficient to unambiguously attribute the droppings to MLGs. We were thus quite confident regarding the reliability of our final genetic dataset.

Our estimate of 79 [68–92] individuals was 1.5 larger than the 57 individuals estimated from the lek counts in 2014. Underestimation of census size from lek counts has already been reported (Aleix-Mata *et al*., 2019, 2024; Jacob *et al*., 2010; Mollet *et al*., 2015) and may result from higher probability of detection of non-territorial males from non-invasive sampling. Indeed, spatial distribution of male around the leks is age-dependent with males 3-years or older defending territories close to lek centers, whereas young males tend to settle in peripheral areas (Pimenta, Morichon and Fons, 2014). Thus, lek counts may underestimate non-territorial males. Extensive studies in related species (especially Sage grouse, *Centrocercus urophasianus*) convincingly demonstrated that all males do not attend leks all days, and, thus, that single counts at leks are likely to underestimate census size (Reese & Bowyer, 2007). Moreover, females only visit the leks for a few days to mate, after which they return to their nesting habitat, usually around 300 m from the lek site in the Pyrenees (Ménoni, 1997). Thus, females are likely less detectable than males during conventional lek counts from tents.

Using CAPWIRE for overall population size (N) in the range of 100 individuals, a mean frequency of observation of 2.89, as observed in our study, generally provides estimates within 10% from the true N and CIs of width < 1/3N. Heterogeneity in capture probability between individuals has been shown to further reduce bias and narrow CIs (Miller *et al*., 2005). Our study thus provided a first reliable estimate of capercaillie population size in the Catalan Nature Reserves. This baseline allows managers to adapt the monitoring scheme to achieve the desired level of accuracy, whether for estimate of census size or assessing the direction of population demographic trend.

Our finding of a balanced sex ratio was rather unexpected, as declining populations often exhibit a male-biased sex ratio. This pattern has been observed in *Tetrao urogallus* populations in the Vosges (70 males and 59 females; Cayuela *et al*., 2019), Central Switzerland (77 males and 46 females; Mollet *et al*., 2015), and *Tetrao urogallus subsp. cantabricus* in the Cantabrian Mountains (32 males and 19 females; Morán-Luis *et al*., 2014), with males generally showing higher average recapture rates. However, the mechanisms driving this relationship remain unclear (but see Donald, 2007).

### The presence of inbreeding suggests a lack of gene flow with neighboring populations

EBVs estimates revealed low genetic diversity and evidence of inbreeding. Our findings indicate that all individuals belong to the same gene pool, and the absence of migrants within the study area suggests that the Western Capercaillie population in Catalan Nature Reserves may be genetically isolated from neighboring populations, resulting in limited gene flow. The inbreeding coefficient at the population level (*H*_O_ = 0.279; *H*_E_ = 0.305; *G*_IS_ = 0.085) was found to be higher in the Catalan Nature Reserves compared to estimates obtained through microsatellite genotyping of other *Tetrao urogallus* populations across their entire European range (Segelbacher *et al*., 2003). However, a similar pattern of inbreeding was observed using SNP genotyping in 24 individuals from the Pyrenees Mountains (mean *F_IS_* = 0.04, range: 0– 0.11). In contrast, higher inbreeding levels were detected in 14 individuals from the endangered *T. u. cantabricus* population inhabiting the Cantabrian Mountains (mean *F_IS_* = 0.10, range: 0– 0.34; Escoda *et al.,* 2023).

The low genetic diversity observed in the Eastern Pyrenees population is consistent with low genetic connectivity to neighboring populations and habitat fragmentation at large scale, owing to the occurrences of topographic barriers and discontinuous forest cover, and at smaller scale through human-induced disturbance (settlements and recreational activities). The Segre Valley separates the central and eastern Pyrenees and has been identified as a putative barrier to gene flow causing genetic differentiation between rock ptarmigan populations (*Lagopus muta*; Bech *et al*., 2009, 2013). Capercaillie and rock ptarmigan occupy the same mountain ranges and are both characterized by short natal dispersal distances (1–2 km, median 5–10 km for capercaillie). We would therefore expect that large topographical obstacles, such as the Segre Valley, may constrain dispersal between capercaillie populations in the Pyrenees (Dunn & Braun, 1985; Small & Rusch, 1989). More extensive sampling across a broader geographic range would be necessary to fully test this hypothesis.

Low genetic variability and inbreeding in capercaillie populations may also result from male philopatry (Pimenta, Morichon, and Fons, 2014), delayed access to reproduction and sex-biased dispersal (Höglund & Alatalo, 1995). Higher inbreeding coefficients in males than in females are in line with the theoretical expectation under kin selection. Outside of the breeding season, males typically stay within their natal winter groups (Willebrandt, 1988). Consequently, leks are often composed of calling males from the same winter groups, promoting breeding among close relatives—a pattern observed in the capercaillie population in the Vosges Mountains (Cayuela *et al*., 2019). Female capercaillies are the primary dispersing sex; however, in highly fragmented habitats, their dispersal capacity is reduced, which increases the likelihood of mating between relatives. Small census and effective population size, combined with low levels of gene flow, accelerates the erosion of genetic diversity through genetic drift, and increases the probability that offspring carry potentially deleterious mutations at the homozygous state (Pekkala *et al.,* 2014).

Overall, these findings suggest that the isolated population of Western capercaillie in the Catalan Nature Reserves warrants special conservation attention. Translocation of individuals from neighboring genetically diverse populations may induce genetic rescue and increase genetic variability. Translocations were successfully integrated into the management strategy of the Pyrenean population of rock ptarmigan (Novoa *et al*., 2020) and should be considered for the capercaillie.

### Individual levels of circulating corticosterone metabolites (FCMs)

The observed trends in individual levels of FCMs were contrary to our initial expectations. Based on the findings of Thiel *et al*., (2011), we had anticipated that individual capercaillie would react to disturbance from winter recreational activities by increasing their corticosterone and thus FCM levels. In light of the levels of human activity documented by Nature Reserves technicians, it seems implausible that disturbance levels were not high enough to induce the production of FCMs by capercaillie. It is possible that the individuals have become accustomed to, or even benefit from, human activities, particularly if human presence deters predators (Berger, 2007; Moss *et al.,* 2014). In previous studies the overall levels of FCMs in grouse were found to be lower than those observed in our study (Thiel *et al*., 2008, 2011), with a notable difference between the sexes (Thiel *et al*., 2008). However, the lowest levels observed in our results were within the range of those previously reported, while others were slightly higher (though they did not reach the maximum levels observed after an ACTH challenge; Thiel et al., 2005).

Our model incorporated few explanatory variables and explained little variance, which suggests that other variables, such as predation pressure, mate finding or abiotic factors, may play a role. However, estimating predator and capercaillie densities either qualitatively or quantitatively is notoriously challenging. Appropriate design would require a comparison between sites, including control sites in which relevant factors are estimated. Additionally, longitudinal studies involving the same animals in different situations would provide valuable insights. Dietary factors, in particular, have the potential to influence FCM levels (Palme, 2019), making them an intriguing avenue for further investigation.

### Variation in individual diet between study sites

Metabarcoding analyses has enabled the detection of 26 plant OTUs that are likely to compose Western Capercaillie diet in the Eastern Pyrenean Mountains. Pinaceae were found in most feces, which confirmed previous knowledge on capercaillie feeding behavior during winter and spring, but we also observed structuration of diet composition at the scale of few kilometers within Catalan Nature Reserves. This pattern may result from differences between study sites in altitude and slope, two forces shaping plant communities. In particular, plants of the Betulaceae family, and plants of the genus *Vaccinium*, possibly the bilberry (*Vaccinium myrtillus*) were almost exclusively present in Py and Mantet sites respectively. Our findings are in line with recent studies based on metabarcoding, which identified 122 plant species from capercaillie feces, and showed that the number of consumed plant species differed between European populations and seasonally between autumn and spring (Chua *et al.,* 2021).

At the individual level, most droppings contents indicate a feeding behavior upon only one dietary resource (*Pinus sp.)* while few drops contained several, yet at most 6 OTUs. Most plants other than Pinaceae that were identified in this study, including *Vaccinium sp.*, *Quercus sp.*, or *Cupressus sp.* were already identified as being part of Capercaillie diet and in particular in neighboring populations in the Pyrenees or in the Cantabrian mountains (Blanco-Fontao *et al.,* 2010; Chua *et al.,* 2021). It would be interesting to know whether the different birds established locally in the Nature Catalan Reserves that make up the same genetic entity behave as different ecological populations using specific trophic niches over time, or whether the birds exhibit opportunistic feeding behavior depending on where they are found. This could be studied by characterizing the diet of birds for which several droppings could be collected from different areas. Unfortunately, this could not be done in this study most likely because of the rather poor quality of plant DNA extracted from droppings. Specific plant DNA extraction kit should be considered to improve plant DNA yields and success of metabarcoding approach.

Diet did not differ between males and females, although females were expected to select more energy-rich resources than males to compensate for resources allocated into egg production (Storch 1993). Sex-based difference in diet selection was detected at local scale, but not at European scale, with females largely preferring *Alnus* compared to males (Chua *et al.,* 2021). The only OTU assigned to *Betulaceae* (possibly *Alnus sp.*) was detected in feces droppings from one male and two females at study site Py, suggesting that *Betulaceae* may be rare habitats used by capercaillie in the Catalan Nature Reserves.

## CONCLUSION

The relationship between species and their habitats significantly influences the estimation of abundance and the effectiveness of monitoring programs for target species, primarily due to the challenge of imperfect detectability. This may in turn increase the risk of inaccurate assessment of the threat status in rare and elusive species and hinder adequate decision-making in management and conservation programs. The use of molecular techniques to assess the ecological and evolutionary status of populations of endangered species can offer interesting complementary data for monitoring conservation actions, bridging the gap between biased field observations and the need for accurate population estimates (Royle *et al.,* 2005; Schwartz *et al.,* 2007). In the present study, we assessed various population parameters, including population census size, sex ratio, genetic structure, genetic diversity, inbreeding levels, FCM levels, and diet composition, thereby enhancing our understanding of the ecology of the Western Capercaillie in the Catalan Nature Reserves. This approach may be applied in neighboring capercaillie populations to estimate current levels of genetic connectivity at the scale of the Pyrenean Mountain range and propose adequate management strategies for the species, including individual translocation if necessary.

## SUPPLEMENTARY FILES

**Appendix 1 :**
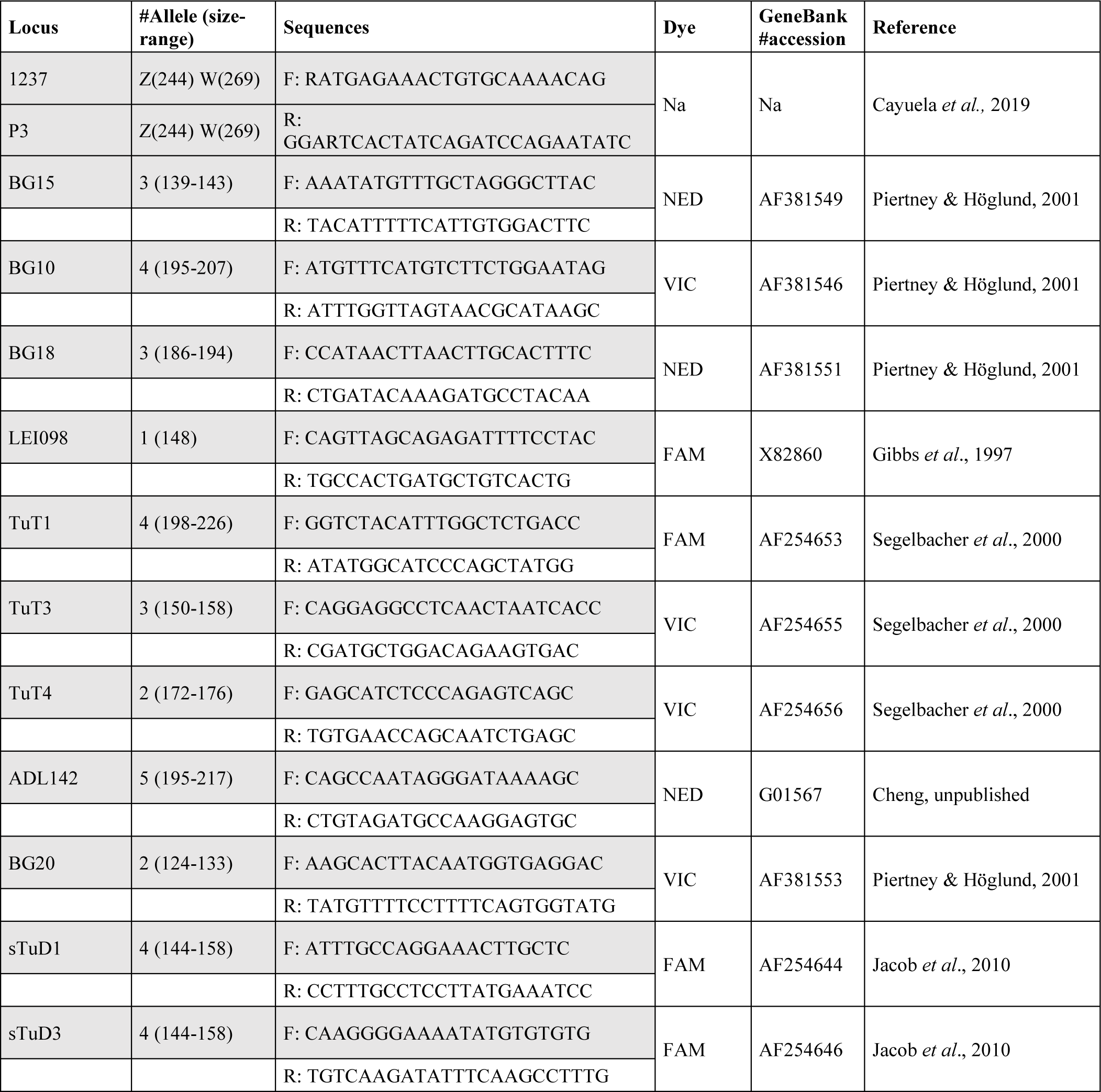
Eleven microsatellites and a *CHD*-gene fragment that were amplified for sexing and genotyping. We indicate locus name, number of alleles, size and range, forward and reverse sequences, fluorescent dye (Blue: FAM, Yellow: VIC, Red: PET, Green: NED), GenBank accession number and reference.

**Appendix 2 :**
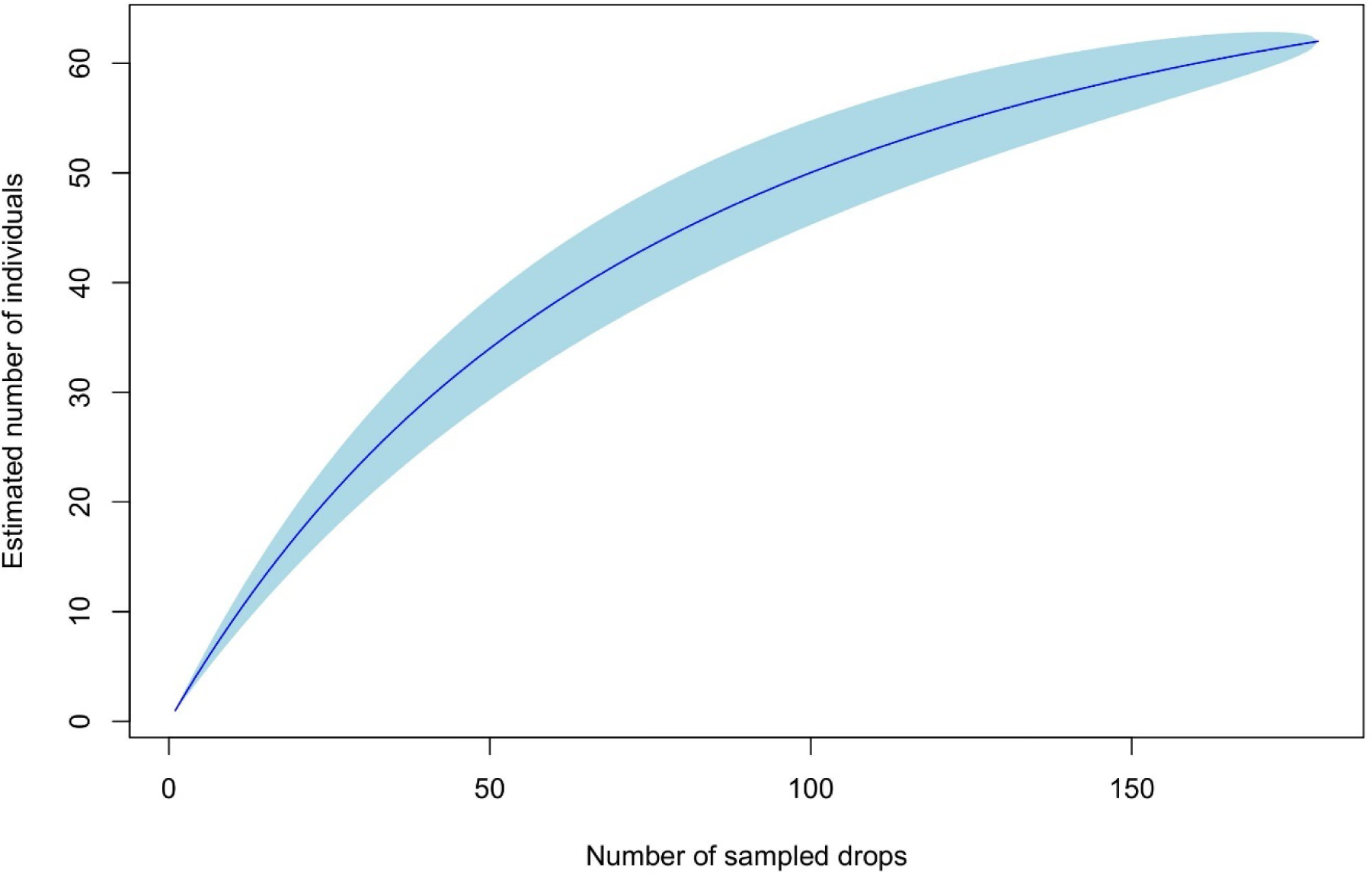
Rarefaction curve which shows the estimated number of individuals according to the number of drops sampled. This curve aims to reach a plateau illustrating the minimum number of drops sampled necessary to obtain all individuals present in the reserve.

**Appendix 3:**
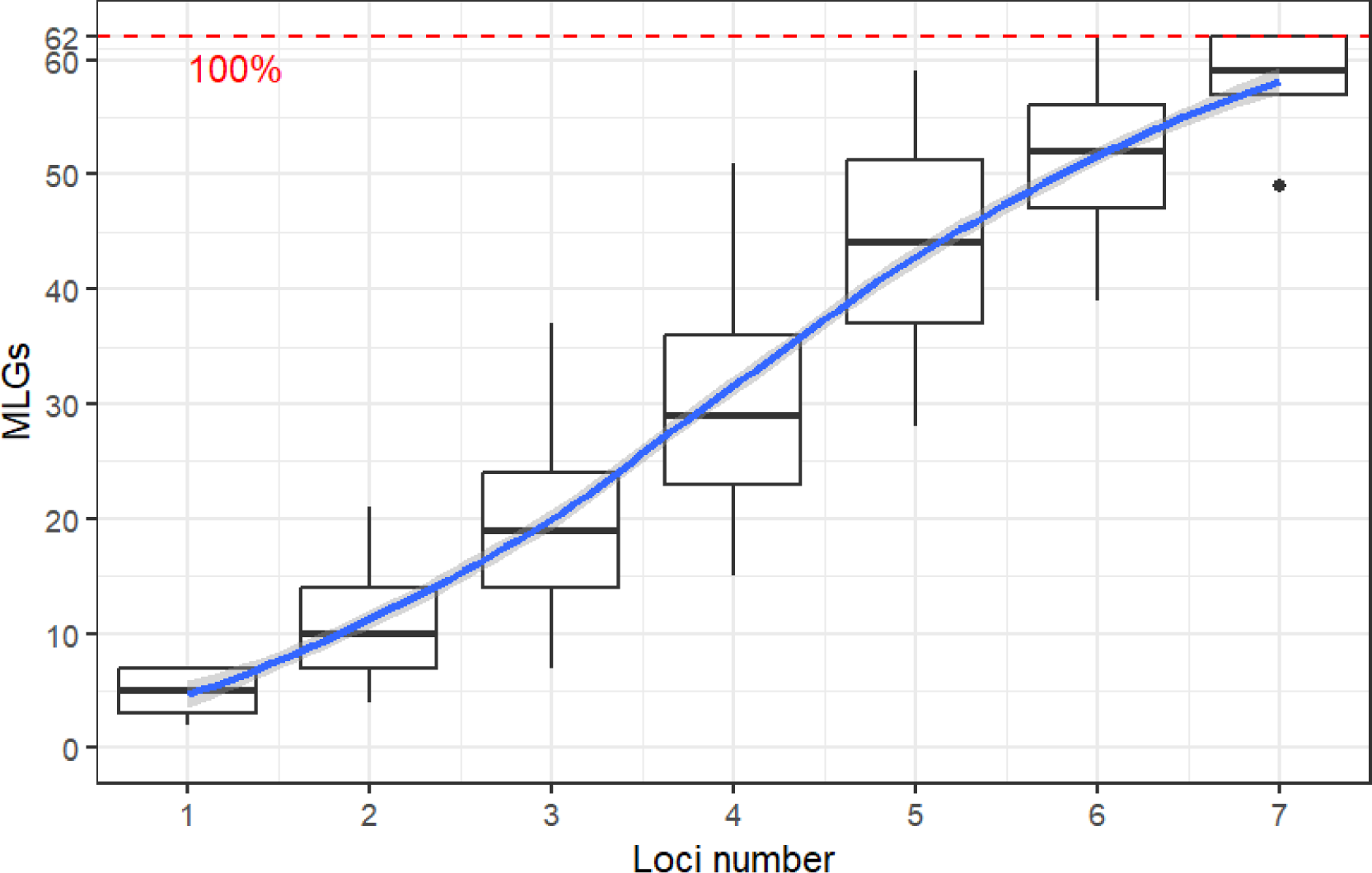
Ability of microsatellite markers. The graphical representation shows that 7 loci are sufficient to obtain 100% of the MLGs within Western Capercaillie population (*Tetrao urogallus subsp. aquitanicus*).

**Appendix 4 :**
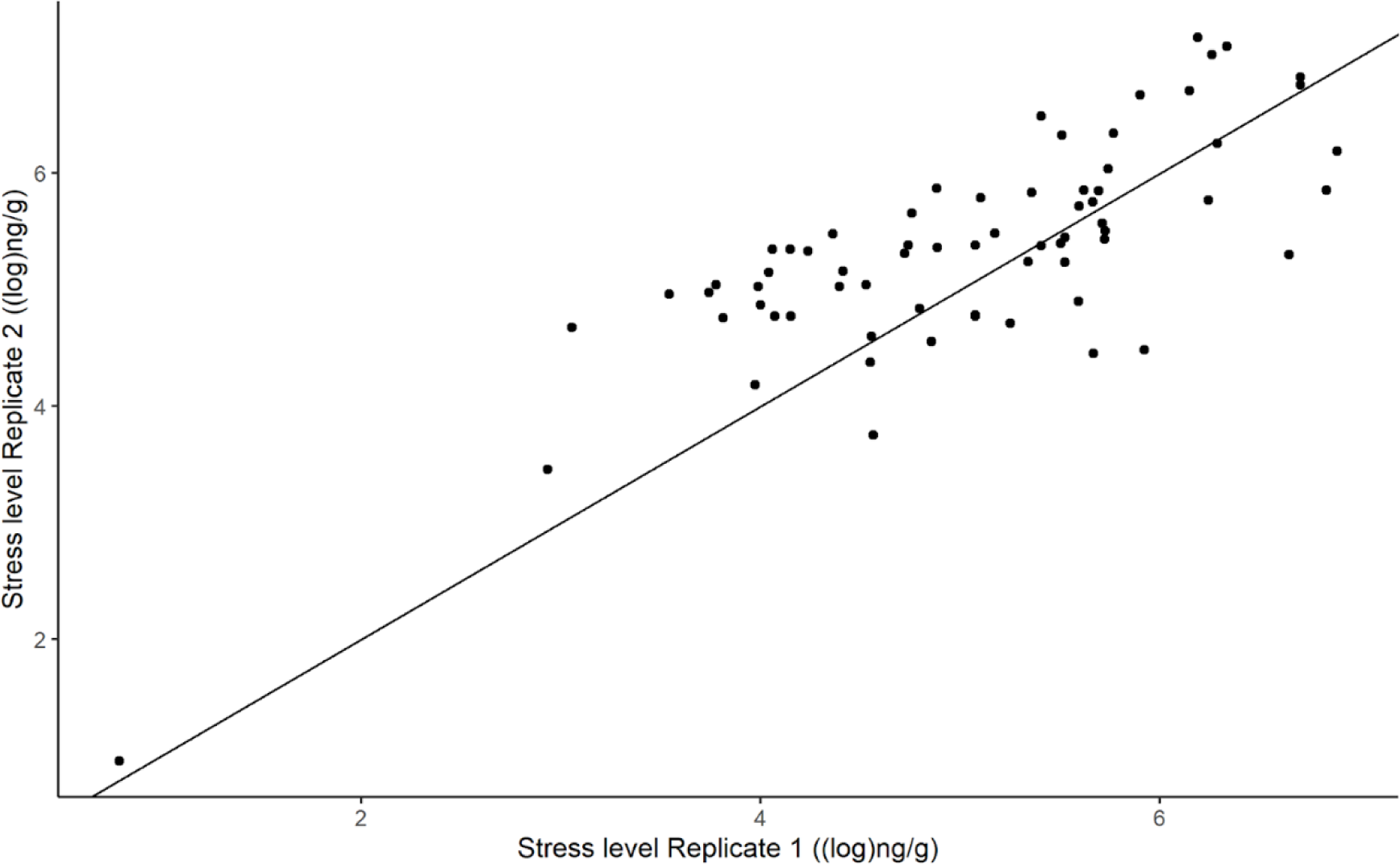
Correlation between Replicate 1 & 2 values. (R² = 0.771, cor.test : p-value = 1.374e-14, t = 9.851, df = 66).

